# CDCP: a visualization and analyzing platform for single-cell datasets

**DOI:** 10.1101/2021.08.24.457455

**Authors:** Yuejiao Li, Tao Yang, Tingting Lai, Lijin You, Fan Yang, Jiaying Qiu, Lina Wang, Wensi Du, Cong Hua, Zhicheng Xu, Jia Cai, Zhiyong Li, Yiqun Liu, Ling Li, Minwen Zhang, Jing Chen, Lei Zhang, Dongsheng Chen, Shiping Liu, Liang Wu, Wenjun Zeng, Bo Wang, Xiaofeng Wei, Longqi Liu, Fengzhen Chen

**Author notes:** These authors contributed equally (Yuejiao Li, Tao Yang, Tingting Lai, Lijin You, Fan Yang, Jiaying Qiu). Correspondence should be addressed to Xiaofeng Wei, Longqi Liu, and Fengzhen Chen.

## Abstract

Advances in single-cell sequencing technology provide a unique approach to characterize the heterogeneity and distinctive functional states at single-cell resolution, leading to rapid accumulation of large-scale single-cell datasets. A big challenge undertaken by research community especially bench scientists is how to simplify the way of retrieving, processing and analyzing the huge number of datasets. Towards this end, we developed Cell-omics Data Coordinate Platform (CDCP), a platform that aims to share and integrate comprehensive single-cell datasets, and to provide a network analysis toolkit for personalized analysis. CDCP contains single-cell RNA-seq and ATAC-seq datasets of 474 572 cells from 6 459 samples in species covering humans, non-human primate models and other animals. It allows querying and visualization of interested datasets and the expression profile of distinct genes in different cell clusters and cell types. Besides, this platform provides an analysis pipeline for non-bioinformatician experimental scientists to address questions not focused by the submitters of the datasets. In summary, CDCP provides a user-friendly interface for researchers to explore, visualize, analyze, download and submit published single-cell datasets and it will be a valuable resource for investigators to explore the global transcriptome profiling at single-cell level.

## 1. Introduction

Recently, the establishment and rapid development of single-cell sequencing technology including sequencing of single-cell genomics, transcriptomes, epigenetics, and spatial transcriptomes makes it possible for researchers to investigate cell heterogeneity, as well as differences in gene expression, epigenetic modifications and spatial information of gene expression at single-cell resolution, which results in the continuous accumulation of single-cell datasets (Clark et al., 2016; Gawad et al., 2016; Longo et al., 2021). However, the lack of a platform for comparative analysis of single-cell datasets is a major obstacle for in-depth mining of existing data.

As the rapid accumulation of single-cell datasets, several databases that integrating datasets in distinct aspects are emerging. For example, CancerSEA was built to explore distinct functional states of cancer cells at single-cell level, covering single-cell transcriptomics of 41 900 cancer single cells from 25 cancer types (Yuan et al., 2019). Abugessaisa et al. established SCPortalen, a database that includes publicly available single-cell datasets from human and mouse (Abugessaisa et al., 2018). Despite their immense efforts, a comprehensive platform that covers single-cell datasets in diverse cell types from all species remains to be established. With the rapid development and accumulation of single-cell datasets, it has brought challenges for the research community, especially the bench scientists, to process massive datasets. Considering the cost of time in learning bioinformatics tools, it is necessary to establish a user-friendly platform for datasets analysis to satisfy the personalized needs of bench researchers.

Here, we developed Cell-omics Data Coordinate Platform (CDCP) to share and integrate complex single-cell datasets. Compared with the aforementioned databases, CDCP contains datasets in a huge number of species including animals, plants and microbes. It allows visualization of each single-cell databases with tSNE cell dimensionality reduction graph, graph of cluster analysis of different cell types, and histogram displaying the number of different cell types. Furthermore, the expression patterns of one or multiple genes in different cells types or clusters can be displayed with cluster graphs and violin graphs, promoting researchers to access and explore the published single-cell datasets. Moreover, CDCP contains detailed information of the biological samples, protocols and library construction methods of the datasets and allows download of the raw sequence and expression matrix of each single-cell dataset. A user-friendly analysis pipeline was developed in the platform to facilitate the reanalysis of single-cell datasets of interest. By providing a single cell expression matrix, users can calculate quality control metrics, annotate highly variable genes, perform principal component analysis, compute a neighborhood graph of observations, embed the neighborhood graph and find marker genes for characterizing groups.

In order to integrate the single-cell datasets, we designed a platform that can expand the database scheme to adapt to different data types generated by single-cell experiments, and developed a web-based user interface named CDCP, which can be freely accessible through https://db.cngb.org/cdcp/.

## 2. Results

### 2.1 Database contents and statistics

CDCP is built to provide a platform for sharing single cell datasets and allow users to query the expression of distinct genes in specific cell types or clusters. At the time of this publication, CDCP contains transcriptomics profiles of 6 467 samples and 474 573 cells from publicly available datasets in species including animals, plants and microbes. Although the existing datasets are collected from only 21 studies, the application of this platform will attract a large number of researchers to upload and share their datasets on it.

CDCP is established and maintained using the Django web framework (https://www.djangoproject.com/), the Python programming language and Pycharm (https://www.jetbrains.com/pycharm/). The Centos7 (https://www.centos.org/) is chosen as the operating system of our server. In terms of system architecture, we use NGINX (http://nginx.org/) to provide static resource access, uWSGI (https://uwsgi-docs.readthedocs.io/en/latest/) to deploy query and download services, Postgresql to store metadata. In terms of data security, the database is deployed in the CNGB (Wang et al., 2019) which has passed the three-level review of information security level protection and the protection capability review of trusted cloud service. Moreover, all services have been deployed with high availability. A series of user-friendly interfaces, including interactive graphs and tables, are used to support its diverse functions. CDCP provides five main functions: exploring data, associated databases, bio-informatics analysis, data visualization and data submission. The user-interface is divided into sub-views as explained below.

### 2.2 Data accessibility

CDCP’s public user interface allows intuitive browsing and searching of available single-cell datasets in the platform. In the “home” page, users can quickly have a view of the statistic of projects, samples, organisms and cells covered in CDCP (Figure 1A). By clicking the buttons of “Project”, “Sample”, “Organism” or “Cell”, users can be directed to the “Explore” page for data retrieval. The summary of datasets in diverse organisms is presented below. Users can get the number of projects, samples, cells, as well as the organs provided in different categories (Figure S1A). Besides, short introduction of shared databases including MBA, NHPCA, HCL and VThunter is provided for users who are interested in (Figure S1A).

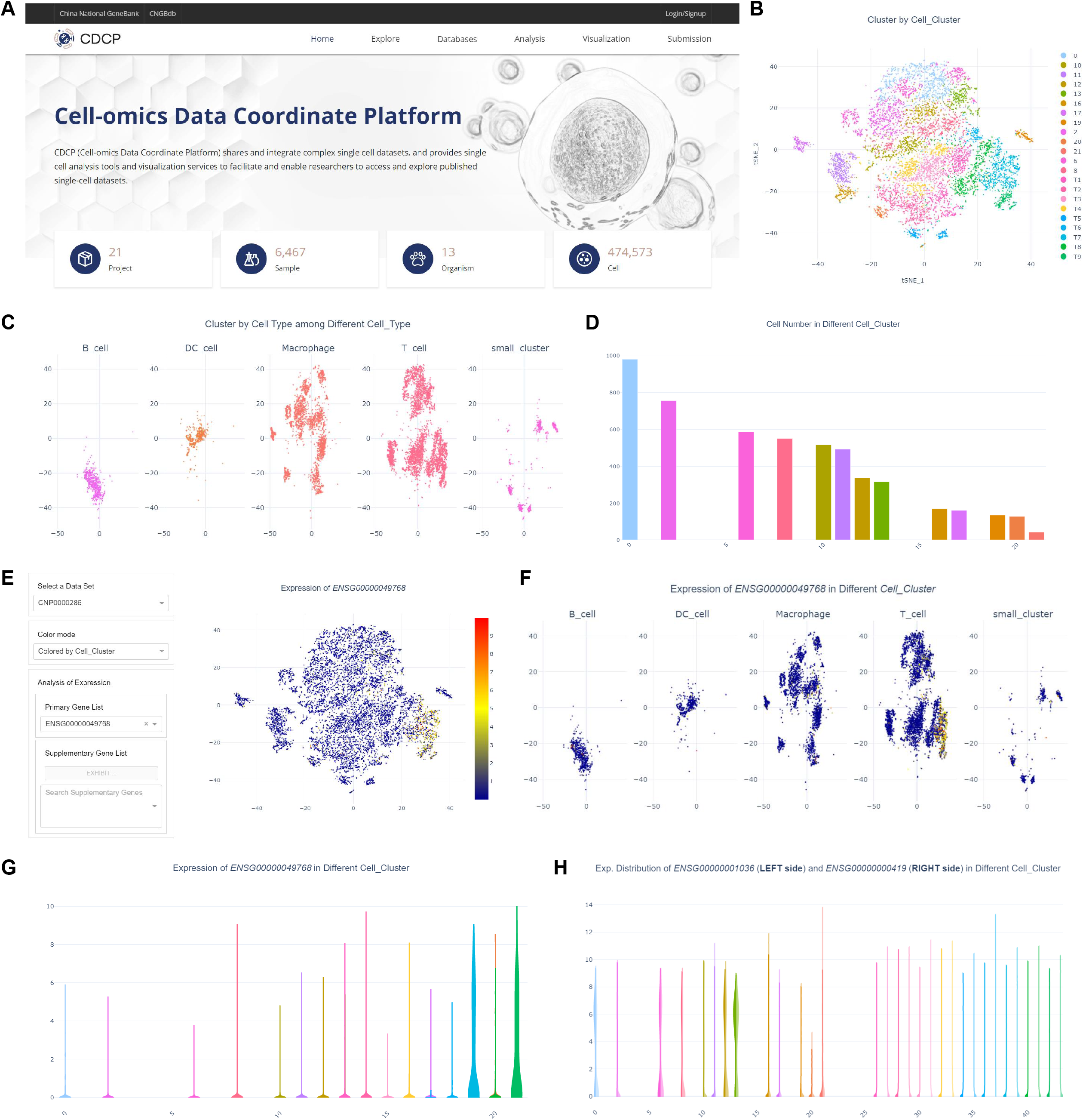

#### 2.2.1 Datasets exploration

In the “Explore” page, one is able to retrieve a single-cell or a group of cells by selecting the attributes of Project, Organism, Tissue, Disease, Library strategy or Release date. The query results that satisfied the search criteria will be displayed including details of the datasets (Figure S1B). Actions for download of raw data and visualization of results are also available. When clicking a specific project ID, users can be directed to the corresponding page showing the detailed meta information about each single-cell sample. This includes the following information: the project name which summarizes the type of the dataset, the abstract of the associated main article publication, the submitter and affiliation of the dataset and the curated metadata for downloading as a text file. Moreover, statistic information including numbers of samples, experiments and runs is presented. By clicking the number of samples, users are able to access detailed information of each sample including age, sex and origin. Similarly, the details of each experiments such as library strategy and library layout can be obtained when clicking the number of experiments.

#### 2.2.2 Associated single-cell databases

CDCP also provides external links to databases of MBA, NHPCA, HCL and VThunter in the “Databases” module. Macaque Brain Atlas (MBA) depicts a macaque brain atlas and allows searching of gene expressions in diverse brain cell types at single-cell resolution. Non-Human Primate Cell Atlas (NHPCA) is a single cell transcriptomics data resource that provides visualization and preliminary analysis of transcriptomic and forthcoming epigenetic single cell data sampled from non-human primate organs or tissues. Human Cell Landscape (HCL) depicts a basic scheme for the human cell landscape to determine the cell type composition of major human organs using single-cell RNA-seq. VThunter allows single-cell screening of virus receptor expression in the whole animal kingdom and predicting of the host tropism of various virus. These available databases will provide more comprehensive information for users who are interested in field of brain research, human, non-human primate models and virus researches.

#### 2.2.3 Data analysis

In the “Analysis” page, a description of Codeplot, the analyzing tool, and the workflow of data analysis are presented (Figure S1C). To facilitate the analysis and sharing of single-cell transcriptome sequencing datasets, single-cell workspace was implemented to manage relevant datasets and perform comprehensive bioinformatics analysis as well as visualization based on the expression matrix datasets archived in CDCP. When clicking “single-cell workspace”, users can be directed to the Codeplot page, which provides a computing-platform with trusted execution environment for bio-informatics analysis. It incorporates the cutting-edge technologies including object storage, secure gene container, blockchain et al. to ensure the security of data sharing and computation in life science. This single-cell analyzing workflow is developed based on the software scanpy (https://scanpy.readthedocs.io). Users are only required to provide a single cell expression matrix (csv/tsv). Sufficient optional adjustment parameters will be given to support user-defined parameters. Each step produces hdf5 file as input for the next step. We also provide each step as a workflow for users to debug.

#### 2.2.4 Data visualization

Visualization of the selected dataset using t-SNE analysis is displayed in the “Visualization” page with point colors representing different cell clusters or cell types (Figure 1B). Users can browse the cell clusters and cell numbers of each cell type in different projects by selecting a project and a dataset in the corresponding drop-down box presented in the hierarchical navigation menu (Figure 1C-D). In addition, by selecting a gene of interest in drop-down box of “Primary Gene List”, users can analyze the expression pattern of this gene across all clusters and cell types (Figure 1E-G). Moreover, the co-expression patterns of two genes in diverse cell types and clusters can also be obtained (Figure 1H).

#### 2.2.5 Data submission

CDCP allows scientists to submit their own single-cell datasets for sharing with the research community. Five major entities are supported by single cell data submission: project, sample, experiment, run and analysis data. The submission of single-cell datasets is mainly divided into two processes: the submission of original sequencing data and the submission of analysis results. By clicking “The submission of sing-cell data” in the “Submission” page, users will be directed to the submission portal page (Figure S1D). Click “Project” and enter the overall description of the research initiative. Then click “Sample” to fill in the detailed information of each sample. Note that each physically unique specimen should be registered as a single sample with a unique set of attributes. The project and sample information shall be submitted before the submission of data documents. Next, in the “Experiment/Run” module, enter the description of sample-specific sequencing library, instrument and sequencing methods and upload the data files following the instructions. Finally, click “Single cell” on the submission portal page and submit the analyzed data files. The detailed steps for submitting datasets in CDCP is available in the “Submission” page to guide users for data submission.

### 2.3 Example applications

To demonstrate the utility and potential application of CDCP, we queried the immune cell database of breast in human as an example. By selecting “homo sapiens” in the drop-down box of “Organism” and “breast” in the drop-down box of “Tissue”, the project that satisfied the search criteria is presented (Figure S1B). This project depicts the single-cell atlas of 9 683 immune cells from 14 samples in triple-negative breast cancer. By clicking the “Visualization” button on the right, users can be directed to the Visualization page. As visualized by the t-SNE map, 22 cell clusters were obtained. These clusters were annotated as T cells (9 clusters), macrophages (6 clusters), B cells (3 clusters), dendritic cells (2 clusters) and 2 unassigned clusters. T cells were the predominant immune cell population, followed by macrophages, B cells and dendritic cells. Distinct signatures of T cell clusters can be revealed by expression patterns of known T cell functional markers and differentially expressed genes (DEGs) among clusters. For instance, in the 9 clusters of T cells, clusters T7 and T9 exhibited remarkable regulatory T cell (Treg) features for high expression of FOXP3 (ENSG00000049768) and IL2RA (ENSG00000134460) (Figure 1E-G, S2A). The DEGs were also identified among clusters, for example the expression of CD200 (ENSG00000091972) in cluster T8, expression of SOX4 (ENSG00000124766) in cluster T9 and expression of ZNF683 (ENSG00000176083) in T6 (Figure S2B-D). These results suggest that CDCP is a reliable and useful platform for in-depth mining of existing single cell datasets.

## 3. Discussion

With the fast development of experimental techniques to observe single cells, such as single cell RNA-seq and spatial transcriptomics, vast amounts of single-cell datasets are becoming available (Saliba et al., 2014; Wu et al., 2021). One of the aims of this study was to build a platform by collecting and integrating the published single-cell datasets, which may facilitate the easy access of the datasets by biology researchers. We developed CDCP that provides comprehensive curated data of single-cell datasets. As of July 2021, CDCP has collected single-cell databases from 6 467 samples and 474 573 cells, which provides a substantial source for research community to access and analyze the datasets of interest. This platform enables retrieval and visualization of single-cell datasets, as well as the expression profiles of a specific gene in different cell clusters or cell types. Besides, CDCP also allows visualization of the co-expression patterns of two genes for scientists who are interested in the correlation between genes.

CDCP contains single-cell RNA-seq, as well as ATAC-seq in different cells from human, non-human primate models, other animals, plants and microbes. This may help scientists in research fields of animals, plants and microbes to retrieve interested datasets in a more convenient way. Besides providing the comprehensive resource of single-cell datasets, an additional strength of CDCP is the supply of the network analysis toolkit (CodePlot), which facilitates customized data processing with a user-friendly web interface and visualization tools.

Additional work will focus on integrating more data sources. In the near future, we plan to upgrade the database with more available genomic data and extend it to cover more species. Moreover, with the establishment and rapid development of spatial transcriptomics sequencing technology, there is a need to integrate spatial transcriptomics datasets in CDCP in the future. The corresponding visualization and analysis tools should be updated too. We believe that CDCP will be a useful platform for the single-cell research community and may help biology researchers to obtain information from different biological processes.

## Acknowledgments

This work was supported by China National GeneBank (CNGB).

## Data availability

**CRediT authorship contribution statement**

## Conflicts of interest

The authors declare no competing interests.

## Supplementary data

## Notes

https://db.cngb.org/cdcp/

### Competing Interest Statement

The authors have declared no competing interest.

