## Supplementary figures and images for "CDCP: a visualization and analyzing platform for single-cell datasets"

### FigureS1

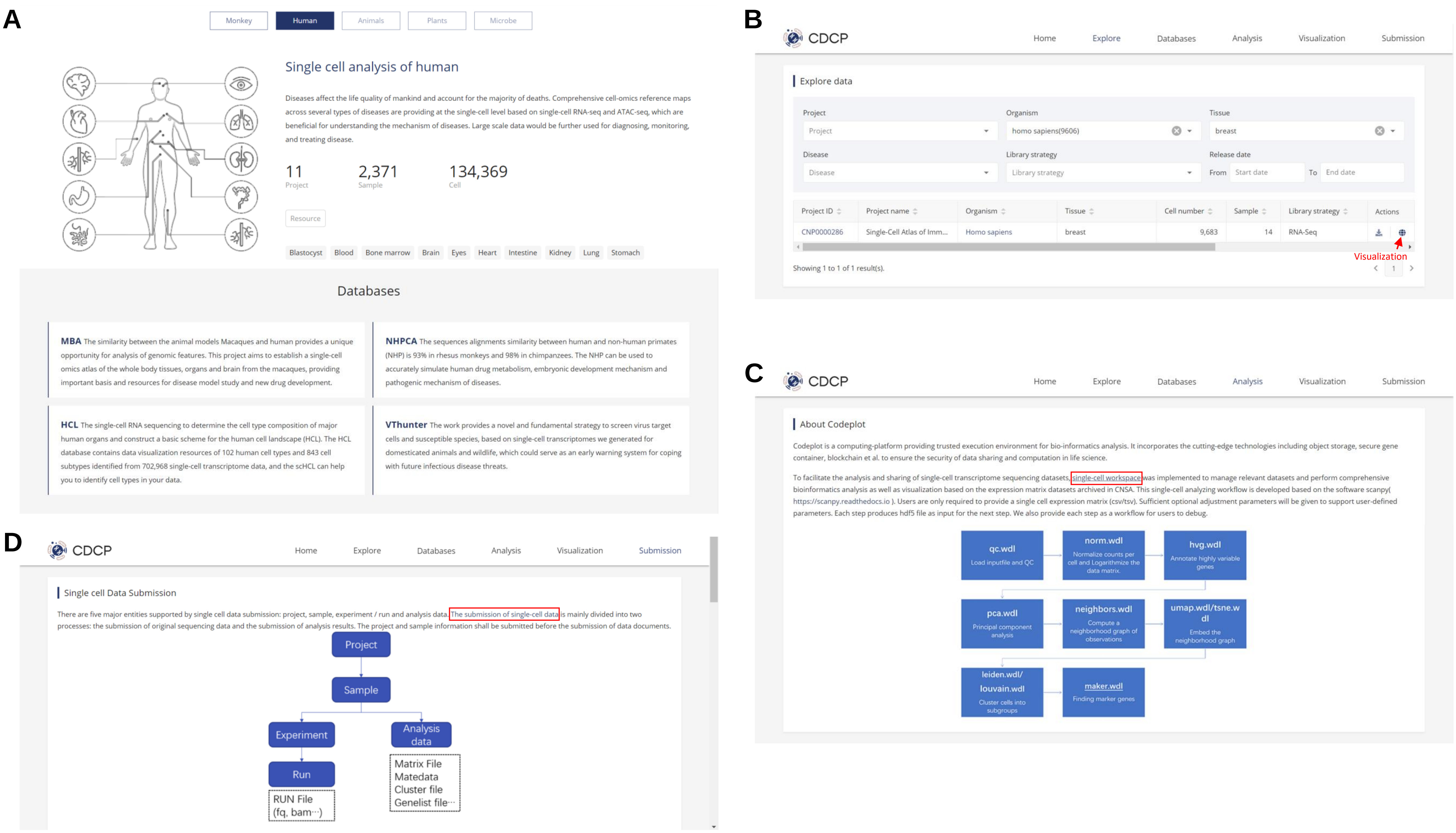
